# Retrospective transcriptome analyses identify *LINC01013* as an activation marker in human dermal fibroblasts

**DOI:** 10.1101/2023.03.21.533694

**Authors:** David M. Dolivo, Adrian E. Rodrigues, Robert D. Galiano, Thomas A. Mustoe, Seok Jong Hong

## Abstract

Study of fibroblast biology, including the process of fibroblast activation, is critical to our understanding of wound healing, tissue fibrosis, and cancer. However, the rapid adoption of next-generation sequencing technologies, particularly single-cell RNA-seq and spatial transcriptomics, has revealed that fibroblast heterogeneity of both healthy and pathological tissues is more complicated than we currently understand. Therefore, a better understanding of molecular players that are not only indicative of but also that contribute to fibroblast activation is critical to piecing together the complete picture and to informing therapeutic strategies to combat associated pathologies. Here we focus on a long-noncoding RNA, *LINC01013*, recently implicated in pathological activation of cardiac fibroblasts and valvular interstitial cell. We analyze several sets of publicly available human transcriptomic data with the aim of determining whether *LINC01013* correlates with fibroblast activation state, and whether compounds that affect fibroblast activation also modulate expression of *LINC01013*. We find that, in numerous independent datasets of healthy and diseased human fibroblasts, *LINC01013* expression is associated with fibroblast activation. We also describe that, even in datasets comprised of small sample sizes, statistically significant correlations exist between expression of *LINC01013* and expression of fibroblast activation markers *ACTA2* and *CCN2*. This finding, while preliminary, suggests that changes in *LINC01013* expression may be an indicator of changes in fibroblast activation state, and that *LINC01013* might functionally contribute to fibroblast activation, lending potential rationale for greater exploration of this lncRNA in the context of tissue fibrosis or tumor stroma.

## Letter Body

Dear Editor,

The paradigm of the activated fibroblast, the myofibroblast, has long been recognized as a critical contributor to dermal fibrosis, as well as to fibrosis in many other tissues. Nevertheless, the advent of transcriptomic technologies, particularly those that can resolve single cells or spatial positioning, has revealed additional layers of complexity to myofibroblast identity, reinforcing the degree to which our understanding of these paradigms has been oversimplified and incomplete^1, 2^. A recent pair of reports identified the long noncoding RNA *LINC01013*, sometimes referred to as *AERRIE*, as a positive regulator of myofibroblast differentiation in human valvular interstitial cells^3^ and human cardiac fibroblasts^4^. These effects are purported to occur through direct potentiation of *CCN2* expression by the lncRNA^3^, and/or through encoding translation of a pro-fibrotic micropeptide^4^. Since the presence and roles of myofibroblasts are somewhat conserved among fibrotic diseases of different tissues^5^, we hypothesized that *LINC01013* might be overexpressed in human dermal myofibroblasts as well. Recently^6^, we generated RNA-seq data of human dermal fibroblasts cultured for 24 hours in the presence or absence of the pro-fibrotic cytokine TGF-β1 and the anti-fibrotic retinoic acid receptor agonist Ch55. We first re-analyzed this data (**Supplementary Methods**) in order to determine how expression of *LINC01013* scaled with promotion and antagonism of dermal fibroblast activation. We first confirmed the pro-fibrotic and anti-fibrotic effects of TGF-β1 and Ch55 by assessing expression of *ACTA2* and *CCN2*, the genes encoding common myofibroblast markers α-SMA and CTGF, respectively (**Fig. S1**). After this verification, we next showed that *LINC01013* was highly upregulated in response to TGF-β1 and highly downregulated upon stimulation with Ch55 (**Fig. 1A**), as would be expected if expression of this gene were correlated with fibroblast activation status. Next we plotted *LINC01013* expression for each RNA-seq sample as a function of *ACTA2* or *CCN2* expression in order to assess whether there was a significant correlation between *LINC01013* and other commonly utilized markers of relative fibroblast activation. Statistical analysis demonstrated a strong, highly significant positive correlation between expression of *LINC01013* and both *ACTA2* and *CCN2* (**Fig. 1B**). In order to explore whether these relationships between *LINC01013* and human dermal fibroblast activation were conserved, rather than an idiosyncrasy characteristic of our own data, we downloaded raw RNA-seq data generated by several other groups who sought to characterize transcriptomes of human dermal fibroblasts in various activation states (**Table S1**). Analyzing this data consistently demonstrated significantly increased expression of *LINC01013* in fibroblasts exposed to TGF-β1 compared to control fibroblasts, in keloid tissue specimens compared to normal skin, and in keloid-derived fibroblasts compared to normal skin-derived fibroblasts (**Fig. 1C**). For each dataset, in spite of the small number of replicates, expression of *LINC01013* across all samples analyzed was found to correlate significantly with expression of *ACTA2, CCN2*, or both, suggesting that *LINC01013* is associated with human dermal fibroblast activation (**Fig. S2-S4**). Notably, treatment with the anti-fibrotic peptide follistatin also reduced expression of *LINC01013*, consistent with a role for *LINC01013* as a marker of fibroblast activation. Thus, we next downloaded RNA-seq data from additional experiments whereby other groups sought to examine transcriptomic effects of anti-fibrotic compounds on human dermal fibroblasts (**Table S1**). Analyzing this data consistently demonstrated decreased expression of *LINC01013* treated with anti-fibrotic compounds FGF-2, JQ1, and nintedanib, while *LINC01013* expression increased upon nintedanib withdrawal (**Fig. 1D**). For each dataset, expression of *LINC01013* across all samples analyzed was found to correlate significantly with expression of *ACTA2, CCN2*, or both (**Fig. S5-S7**). Taken together, these data collectively suggest that *LINC01013* is overexpressed in fibrotic dermal fibroblasts, and that alleviation of fibroblast activation is associated with *LINC01013* downregulation, denoting *LINC01013* as a potential biomarker for pathological fibroblast activation. It is important to note that the scaling of *LINC01013* upwards and downwards with fibroblast activation status is insufficient to conclude that its overexpression is causally involved in fibrotic pathology, but recent reports describing that *LINC01013* drives fibroblast activation in cardiac fibroblasts^4^ and valvular interstitial cells^3^, pathological processes analogous to persistent fibroblast activation characteristic of fibrotic dermal pathologies, suggests that this possibility is worthy of further pursuit and investigation.

**Figure 1.**
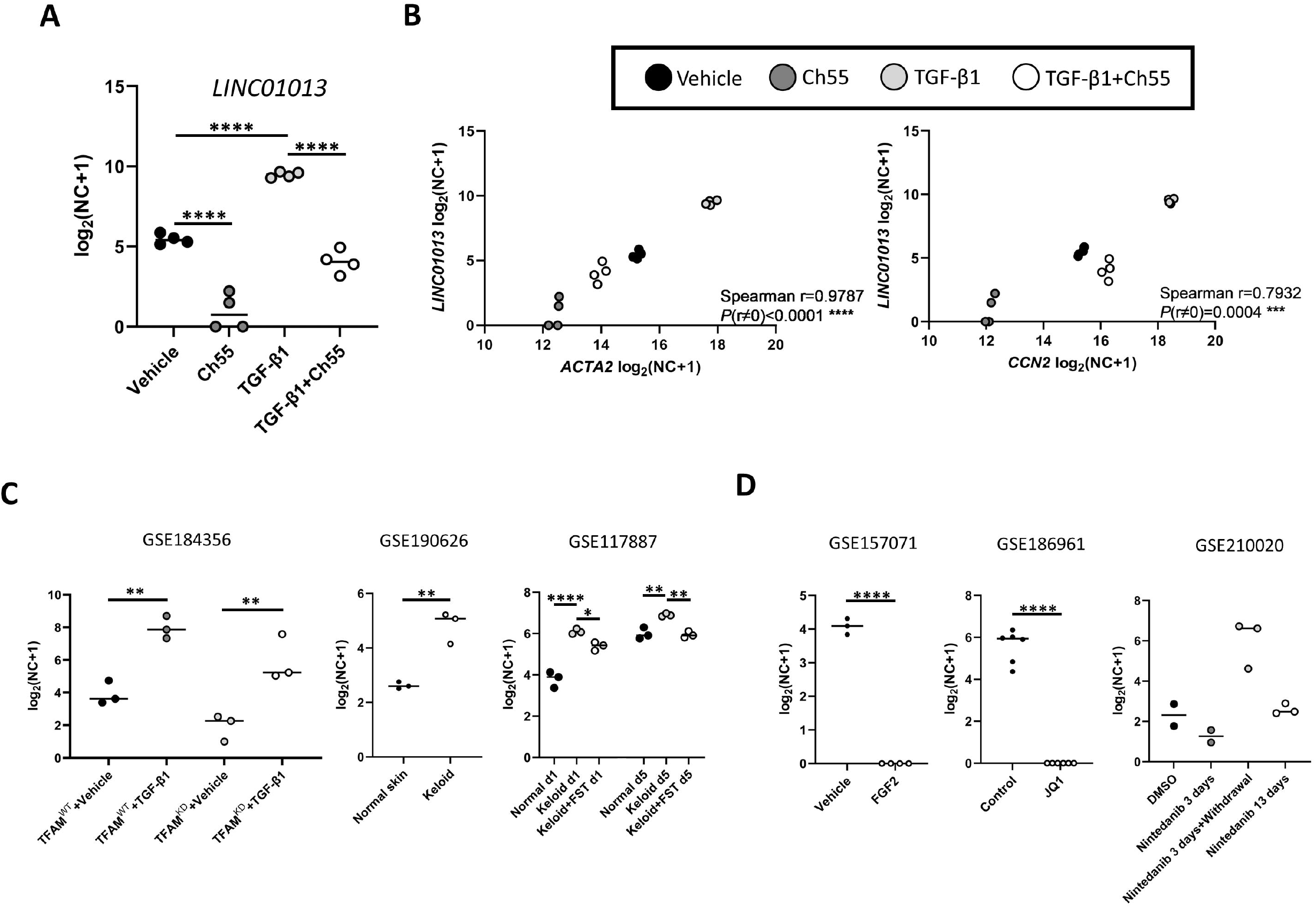
LINC01013 expression scales with human dermal fibroblast activation and correlates with ACTA2 and CCN2 expression. RNA-seq data from human samples were analyzed and normalized counts extracted from various RNA-seq datasets. (A) Plot of normalized expression levels of *LINC01013* in human fibroblasts exposed to Vehicle, Ch55, TGF-β1, or both Ch55 and TGF-β1 for 24 hours. (B) Plot of per-sample *LINC01013* expression as a function of *ACTA2* (left) and *CCN2* (right) expression. (C) Normalized expression of *LINC01013* in human samples representative of fibrotic fibroblast responses. (D) Normalized expression of *LINC01013* in human samples representative of anti-fibrotic fibroblast responses.

## ACKNOWLEDGMENTS

None.

