## Supplementary figures and images for "Retrospective transcriptome analyses identify *LINC01013* as an activation marker in human dermal fibroblasts"

### Fig. S1

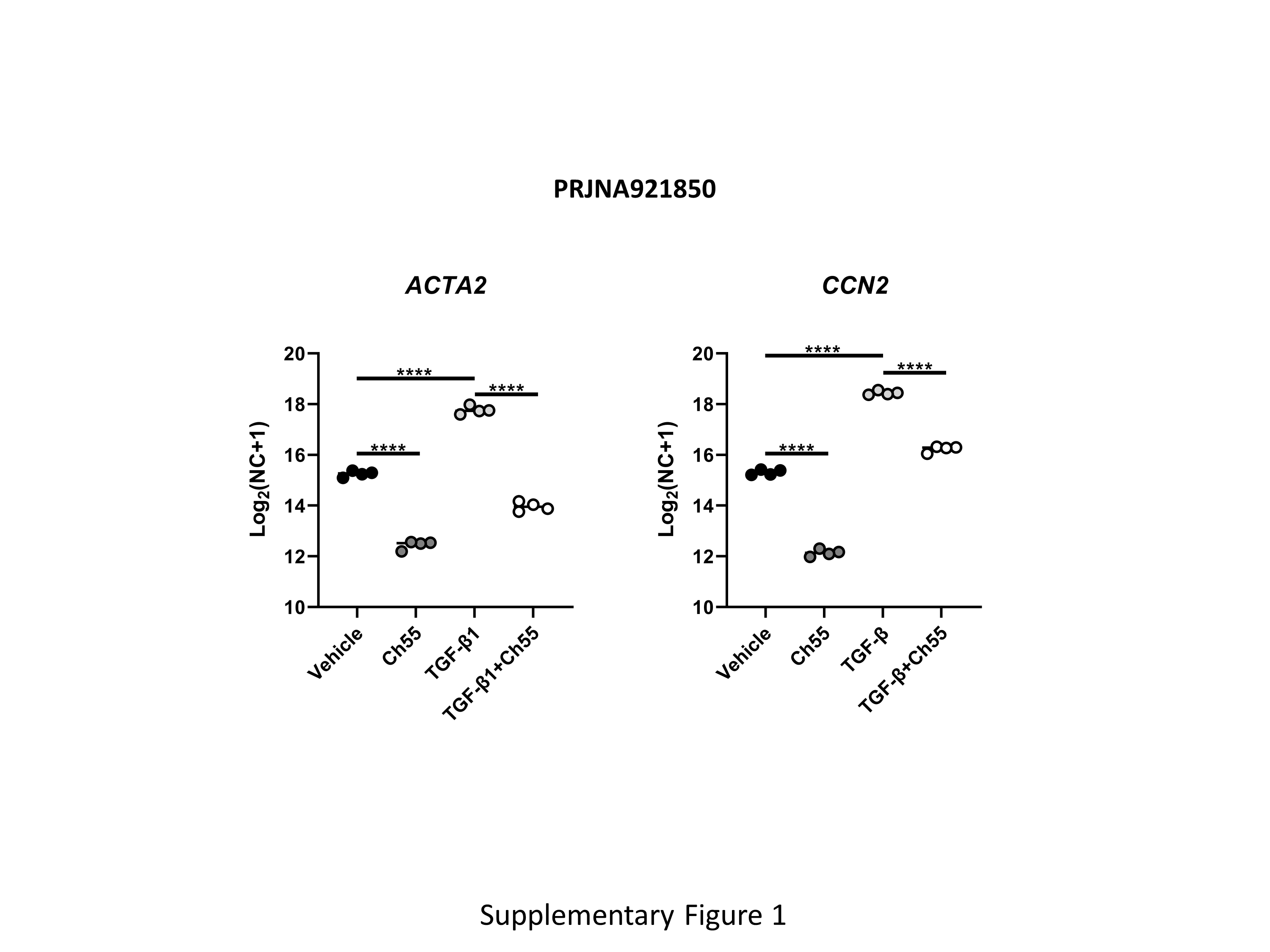

### Fig. S2

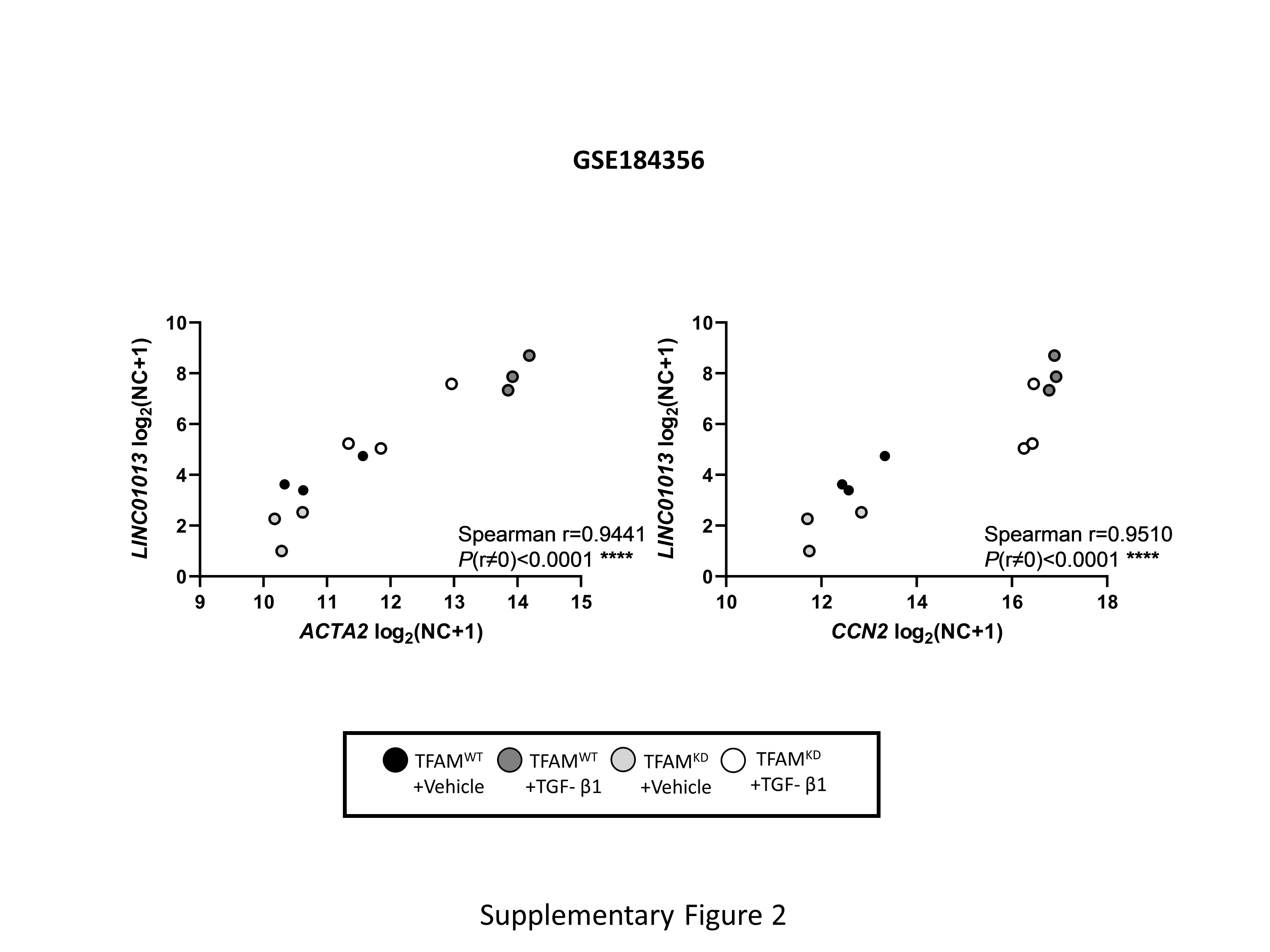

### Fig. S3

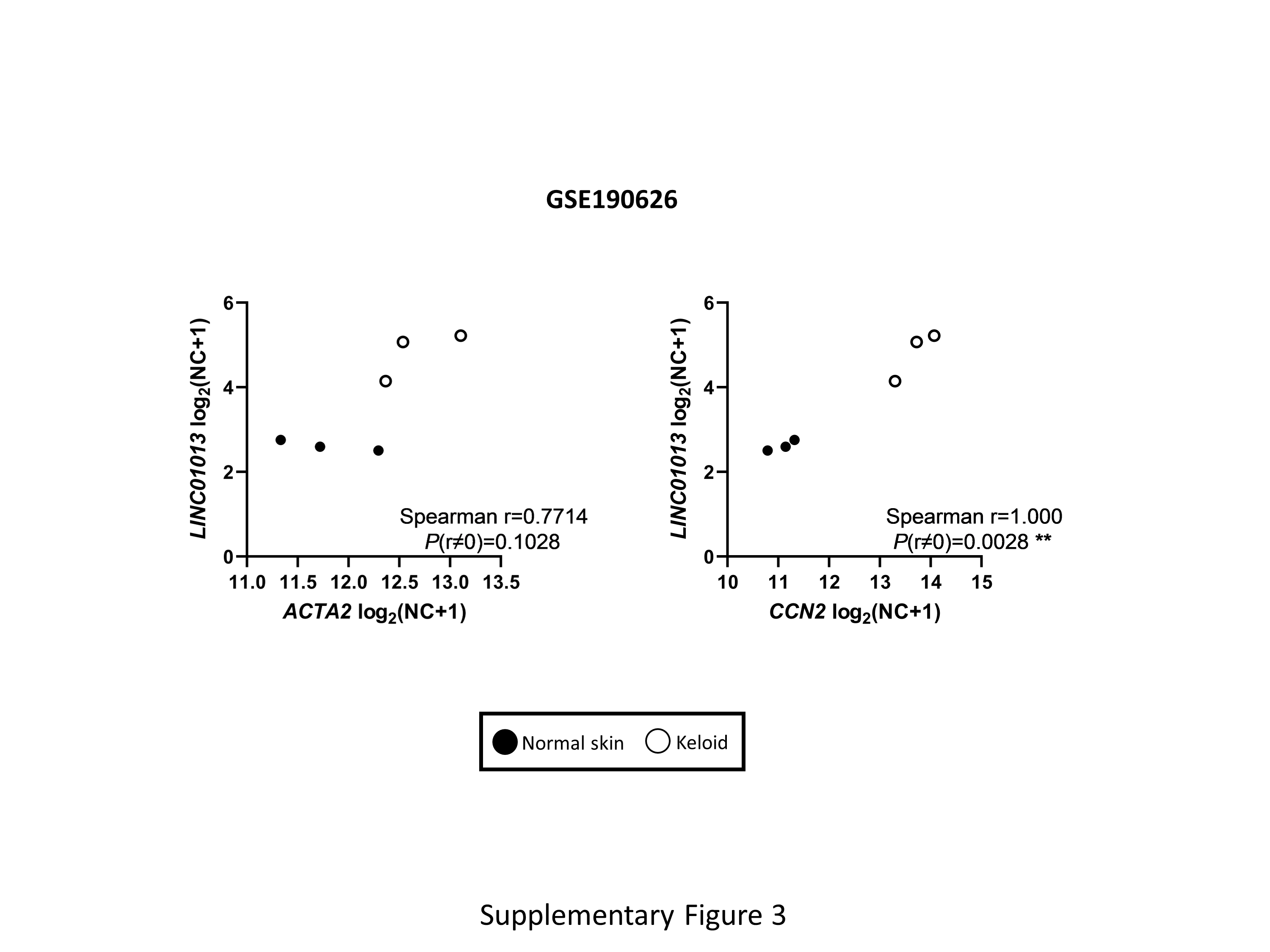

### Fig. S4

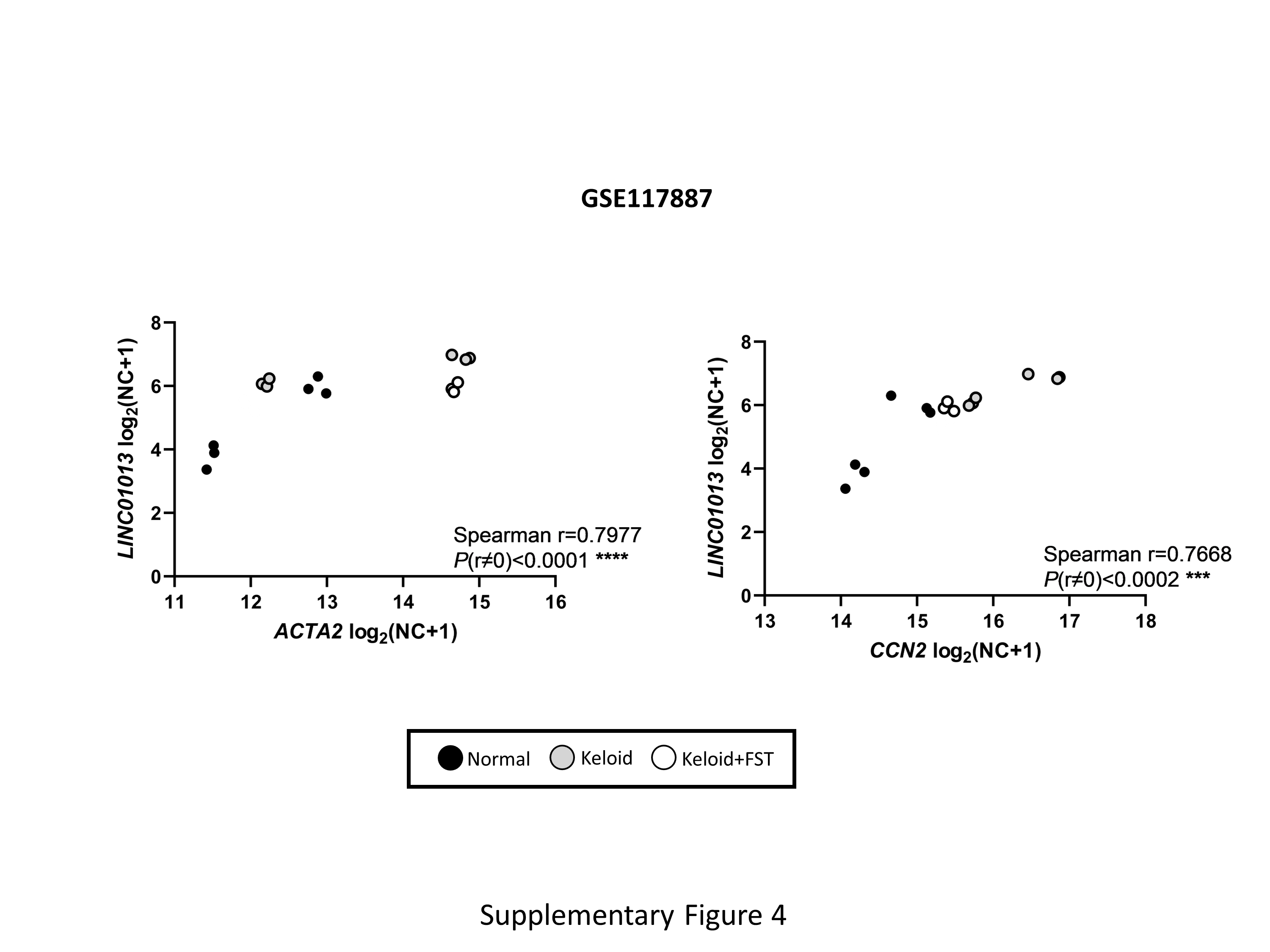

### Fig. S5

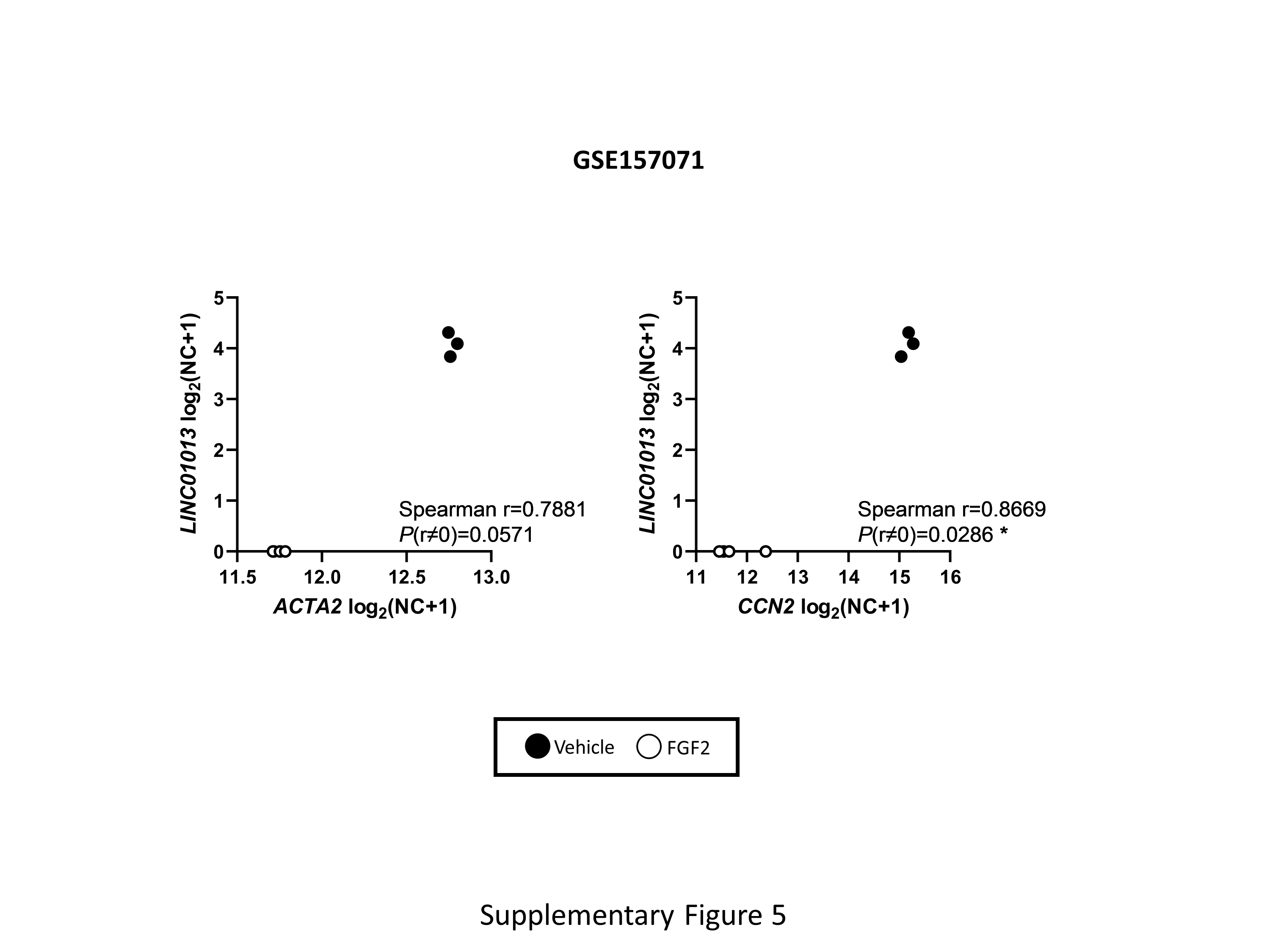

### Fig. S6

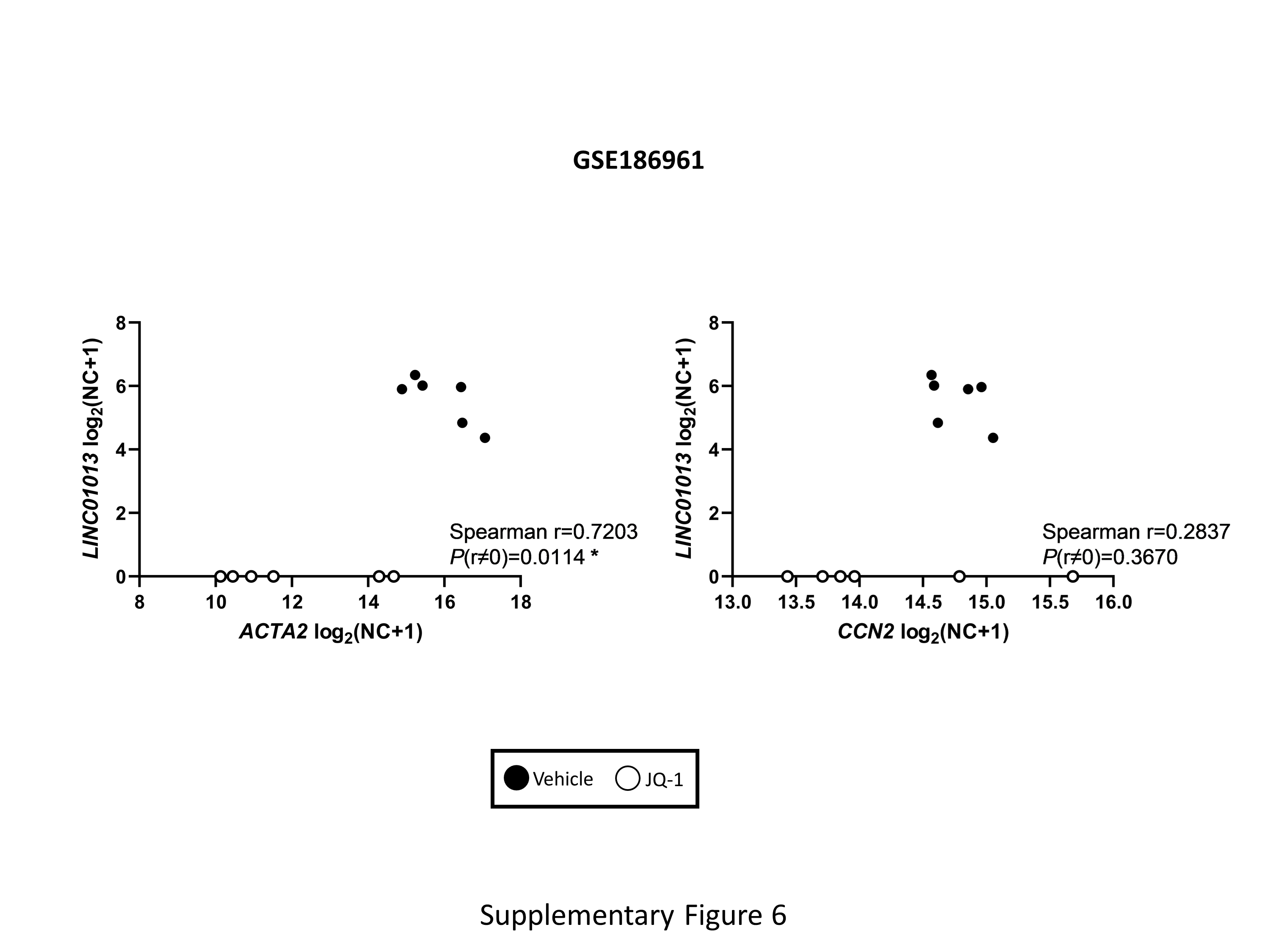

### Fig. S7

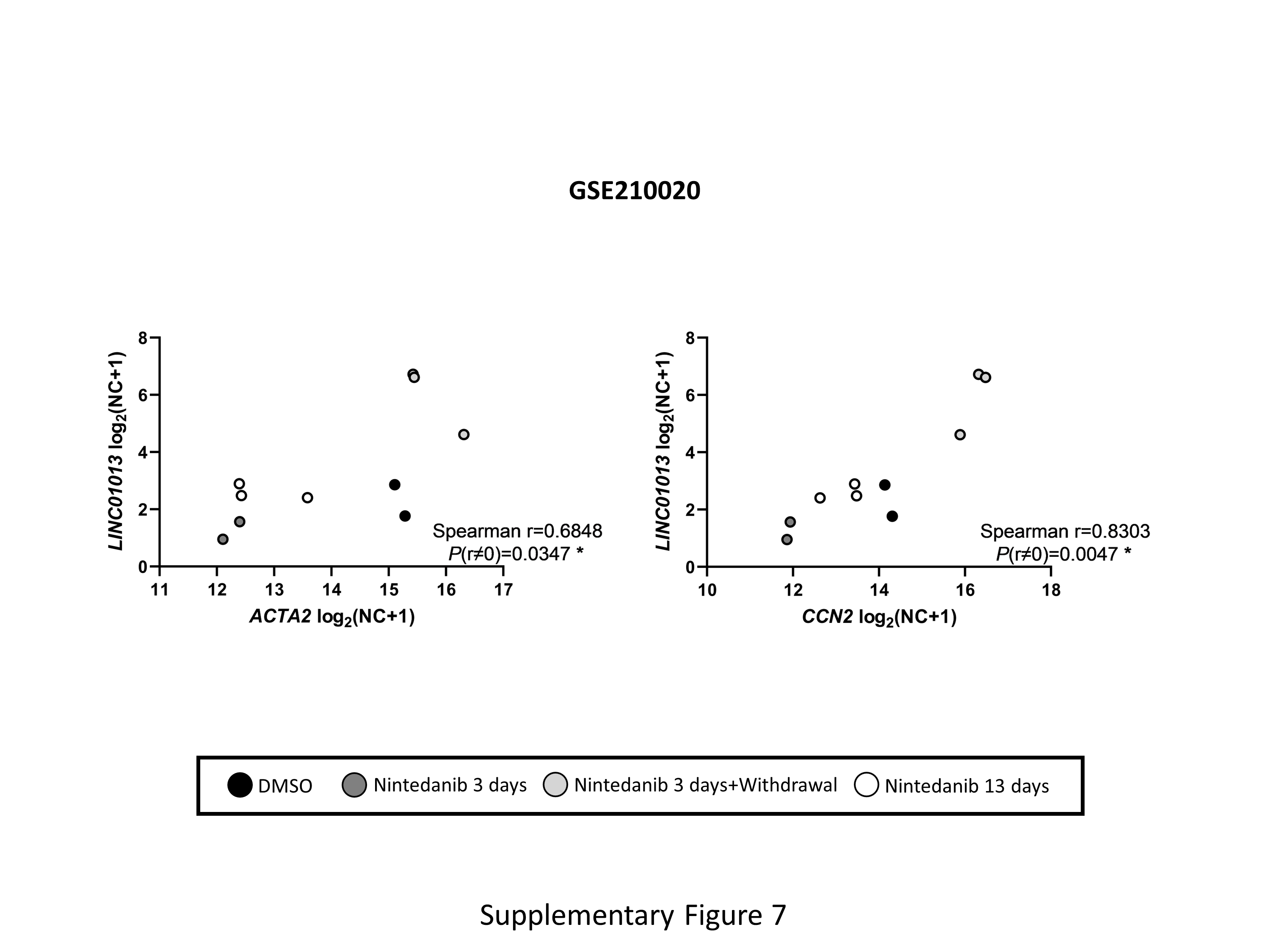
