## Supplementary Figure Legends for "Retrospective transcriptome analyses identify *LINC01013* as an activation marker in human dermal fibroblasts"

*Supplementary Figure 1. ACTA2 and CCN2 dysregulation from treatment with Ch55 and TGF-β1*

Normalized count values of (left) *ACTA2* and (right) *CCN2* in primary human foreskin fibroblasts treated with Ch55 and TGF-β1.

*Supplementary Figure 2. Correlation of LINC01013 with fibroblast activation markers in human dermal fibroblasts in response to TGF-β1*

Normalized count values of *LINC01013* as a function of (left) *ACTA2* and (right) *CCN2* in human dermal fibroblasts infected with control or TFAM^KD^ siRNA, in response to TGF-β1 stimulus.

*Supplementary Figure 3. Correlation of LINC01013 with fibroblast activation markers in human keloid and normal skin tissues*

Normalized count values of *LINC01013* as a function of (left) *ACTA2* and (right) *CCN2* in normal human skin and human keloid tissues.

*Supplementary Figure 4. Correlation of LINC01013 with fibroblast activation markers in human keloid fibroblasts and normal fibroblasts, and upon exposure to follistatin 288*

Normalized count values of *LINC01013* as a function of (left) *ACTA2* and (right) *CCN2* in normal human dermal fibroblasts and human keloid fibroblasts, and in human keloid fibroblasts in response to follistatin 288 treatment.

*Supplementary Figure 5. Correlation of LINC01013 with fibroblast activation markers in human dermal fibroblasts in response to FGF-2*

Normalized count values of *LINC01013* as a function of (left) *ACTA2* and (right) *CCN2* in normal human dermal fibroblasts, in response to FGF-2 treatment.

*Supplementary Figure 6. Correlation of LINC01013 with fibroblast activation markers in human scleroderma fibroblasts in response to JQ1*

Normalized count values of *LINC01013* as a function of (left) *ACTA2* and (right) *CCN2* in primary human dermal fibroblasts isolated from diffuse systemic sclerosis, in response to treatment with JQ1.

*Supplementary Figure 7. Correlation of LINC01013 with fibroblast activation markers in primary human foreskin fibroblasts upon exposure and withdrawal of nintedanib*

Normalized count values of *LINC01013* as a function of (left) *ACTA2* and (right) *CCN2* in primary human foreskin fibroblasts, in response to stimulation and withdrawal of nintedanib.
