## Supplementary Methods for "Retrospective transcriptome analyses identify *LINC01013* as an activation marker in human dermal fibroblasts"

**RNA-seq analysis**

Raw FASTQ sequence data were obtained from the European Nucleotide Archive via direct import into Galaxy^1^. Low-quality sequence files were trimmed using Trimmomatic^2^. Trimmed reads were mapped to the human genome (*hg38*) construction using HISAT2^3^, and mapped reads were assigned to genomic features using featureCounts^4^. Normalized counts output using deseq2^5^ were used for downstream analyses.

**Statistical analysis**

All graphs were constructed and statistical comparisons made using Graphpad Prism (Graphpad, San Diego, CA). All normalized count values were adjusted by addition of exactly 1 in order to enable log transformation. Sample normalized counts are depicted as individual values with median summary bars. Comparisons of expression of single genes between two samples were performed using unpaired, two-tailed Student’s t-tests, and comparisons of gene expression among more than two samples were performed using one-way ANOVAs with Tukey’s post-hoc analysis for pairwise comparisons, with selected comparisons of interest depicted on the respective graphs. For correlational analyses, Spearman r values and *P* values were computed using nonparametric two-tailed analyses. For all comparisons, **P*<0.05,***P*<0.01,****P*<0.001,*****P*<0.0001. Only datasets for which all sample groups contained at least three replicates were subjected to statistical analysis for difference in gene expression. All datasets presented were analyzed for statistical correlation.

**Supplementary Methods References**

1	Jalili V, Afgan E, Gu Q et al. The Galaxy platform for accessible, reproducible and collaborative biomedical analyses: 2020 update. *Nucleic acids research* 2020; **48**(W1): W395-W402.

2	Bolger AM, Lohse M, Usadel B. Trimmomatic: a flexible trimmer for Illumina sequence data. *Bioinformatics* 2014; **30**(15): 2114-2120.

3	Kim D, Paggi JM, Park C, Bennett C, Salzberg SL. Graph-based genome alignment and genotyping with HISAT2 and HISAT-genotype. *Nature biotechnology* 2019; **37**(8): 907-915.

4	Liao Y, Smyth GK, Shi W. featureCounts: an efficient general purpose program for assigning sequence reads to genomic features. *Bioinformatics* 2014; **30**(7): 923-930.

5	Love MI, Huber W, Anders S. Moderated estimation of fold change and dispersion for RNA-seq data with DESeq2. *Genome biology* 2014; **15**(12): 1-21.
