## Supplementary Table for "Retrospective transcriptome analyses identify *LINC01013* as an activation marker in human dermal fibroblasts"

**Supplementary Table 1.** Sources for data used for analysis

| Experimental condition(s) | First author, year  published | Database Accession Number | Cell/tissue source |
| --- | --- | --- | --- |
| 1μM Ch55, 10ng/mL TGF-β1, 24 hours | Dolivo et al. 2023 | PRJNA921850 (NCBI SRA) | Primary human neonatal foreskin fibroblasts |
| 10ng/mL TGF-β1  24 hours | Zhou et al. 2022 | GSE184356  (NCBI GEO) | Primary human dermal fibroblasts transfected with control or *TFAM* siRNA |
| None | Xie et al. 2021 | GSE190626  (NCBI GEO) | Normal skin and keloid specimens |
| 100ng/mL Follistatin 288  1 or 5 days | Ham et al. 2021 | GSE117887  (NCBI GEO) | Primary human dermal fibroblasts from healthy skin or keloid |
| 10ng/mL FGF-2  48 hours | Wu et al. 2021 | GSE157071  ^1^(NCBI GEO) | Primary human dermal fibroblasts |
| 1μM JQ1  48 hours | Vichaikul et al. 2022 | GSE186961  (NCBI GEO) | Primary human fibroblasts from diffuse systemic sclerosis |
| 5μM nintedanib  3 or 13 days | Cho et al. 2022 | GSE210020  (NCBI GEO) | Primary human neonatal foreskin fibroblasts |

Table abbreviations:

TGF-β1: transforming growth factor beta 1

FGF-2: fibroblast growth factor 2

*TFAM*: mitochondrial transcription factor A

siRNA: short inhibitory RNA

NCBI: National Center for Biotechnology Information

SRA: sequence read archive

GEO: gene expression omnibus

**Supplementary Table References**

1	Dolivo DM, Rodrigues AE, Galiano RD, Mustoe TA, Hong SJ. Prediction and demonstration of retinoic acid receptor agonist Ch55 as an anti-fibrotic agent in the dermis. Journal of Investigative Dermatology 2023.

2	Zhou X, Trinh‐Minh T, Tran‐Manh C et al. Impaired Mitochondrial Transcription Factor A Expression Promotes Mitochondrial Damage to Drive Fibroblast Activation and Fibrosis in Systemic Sclerosis. Arthritis & Rheumatology 2022; 74(5): 871-881.

3	Xie J, Chen L, Cao Y et al. Single-cell sequencing analysis and weighted co-expression network analysis based on public databases identified that TNC is a novel biomarker for keloid. Frontiers in Immunology 2021; 12: 783907.

4	Ham S, Harrison C, de Kretser D, Wallace EM, Southwick G, Temple‐Smith P. Potential treatment of keloid pathogenesis with follistatin 288 by blocking the activin molecular pathway. Experimental Dermatology 2021; 30(3): 402-408.

5	Wu B, Tang X, Zhou Z, Ke H, Tang S, Ke R. RNA sequencing analysis of FGF2-responsive transcriptome in skin fibroblasts. PeerJ 2021; 9: e10671.

6	Vichaikul S, Gurrea-Rubio M, Amin MA et al. Inhibition of bromodomain extraterminal histone readers alleviates skin fibrosis in experimental models of scleroderma. JCI insight 2022; 7(9).

7	Cho H-J, Hwang J-A, Yang EJ et al. Nintedanib induces senolytic effect via STAT3 inhibition. Cell death & disease 2022; 13(9): 760.
